# Finding decodable information that can be read out in behaviour

**DOI:** 10.1101/248583

**Authors:** Tijl Grootswagers, Radoslaw M. Cichy, Thomas A. Carlson

## Abstract

Multivariate decoding methods applied to neuroimaging data have become the standard in cognitive neuroscience for unravelling statistical dependencies between brain activation patterns and experimental conditions. The current challenge is to demonstrate that decodable information is in fact used by the brain itself to guide behaviour. Here we demonstrate a promising approach to do so in the context of neural activation during object perception and categorisation behaviour. We first localised decodable information about visual objects in the human brain using a multivariate decoding analysis and a spatially-unbiased searchlight approach. We then related brain activation patterns to behaviour by testing whether the classifier used for decoding can be used to predict behaviour. We show that while there is decodable information about visual category throughout the visual brain, only a subset of those representations predicted categorisation behaviour, which were strongest in anterior ventral temporal cortex. Our results have important implications for the interpretation of neuroimaging studies, highlight the importance of relating decoding results to behaviour, and suggest a suitable methodology towards this aim.

## 1 Introduction

Multivariate pattern analysis (MVPA), also called brain decoding, is a powerful tool to establish statistical dependencies between experimental conditions and brain activation patterns (Carlson, Schrater, & He, 2003; Cox & Savoy, 2003; Haxby et al., 2001; Haynes, 2015; Kamitani & Tong, 2005; Kriegeskorte, Goebel, & Bandettini, 2006). In these analyses, an implicit assumption often made by experimenters is that if information can be decoded, then this information is used by the brain in behaviour (de-Wit, Alexander, Ekroll, & Wagemans, 2016; Ritchie, Kaplan, & Klein, 2017). However, the decoded information could be different (e.g., epiphenomenal) from the signal that is relevant for the brain to use in behaviour (de-Wit et al., 2016; Williams, Dang, & Kanwisher, 2007), highlighting the need to relate decoded information to behaviour. Importantly, this implicit assumption of decoding models leads to testable predictions about task performance (Naselaris, Kay, Nishimoto, & Gallant, 2011). Previous work has for example correlated decoding performances to behavioural accuracies (Bouton et al., 2018; Freud, Culham, Plaut, & Behrmann, 2017; Raizada, Tsao, Liu, & Kuhl, 2010; van Bergen, Ji Ma, Pratte, & Jehee, 2015; Walther, Caddigan, Fei-Fei, & Beck, 2009; Williams et al., 2007). However, this does not model how individual experimental conditions relate to behaviour. Another approach has been to compare neural and behavioural similarity structures (Bracci & Op de Beeck, 2016; Cichy, Kriegeskorte, Jozwik, Bosch, & Charest, 2017; Cohen, Dennett, & Kanwisher, 2016; Grootswagers, Kennedy, Most, & Carlson, 2017; Haushofer, Livingstone, & Kanwisher, 2008; Mur et al., 2013; Proklova, Kaiser, & Peelen, 2016; Wardle, Kriegeskorte, Grootswagers, Khaligh-Razavi, & Carlson, 2016). While this approach allows to link behaviour and brain patterns at the level of single experimental conditions, it is unclear how this link carries over to decision making behaviour such as categorisation (but see Cichy et al., (2017) for recent developments).

Recently, a novel methodological approach, called the distance-to-bound approach (Ritchie & Carlson, 2016), has been proposed to connect brain activity directly to perceptual decision-making behaviour at the level of individual experimental conditions. The rationale behind this approach (Bouton et al., 2018; Carlson, Ritchie, Kriegeskorte, Durvasula, & Ma, 2014; Kiani, Cueva, Reppas, & Newsome, 2014; Philiastides & Sajda, 2006; Ritchie & Carlson, 2016) is that for decision-making tasks, the brain applies a decision boundary to a neural activation space (DiCarlo & Cox, 2007). Similarly, MVPA classifiers fit multi-dimensional hyperplanes to separate a neural activation space. In classic signal-detection theory (Green & Swets, 1966) and evidence-accumulation models of choice behaviour (Brown & Heathcote, 2008; Gold & Shadlen, 2007; Ratcliff & Rouder, 1998; Smith & Ratcliff, 2004), the distance of the input to a decision boundary reflects the ambiguity of the evidence for the decision (Green & Swets, 1966). Decision evidence, in turn, predicts choice behaviour (e.g., Ashby, 2000; Ashby & Maddox, 1994; Britten, Newsome, Shadlen, Celebrini, & Movshon, 1996; Gold & Shadlen, 2007; Shadlen & Kiani, 2013) which also has clear neural correlates (e.g., Britten et al., 1996; Ratcliff, Philiastides, & Sajda, 2009; Roitman & Shadlen, 2002). If for a decision task (e.g., categorisation), the brain uses the same information as the MVPA classifier, then the classifier’s hyperplane reflects the brain’s decision boundary. This in turn predicts that distance to the classifier’s hyperplane negatively correlates with reaction times for the decision task. In the distance-to-bound approach, finding such a negative distance-RT-correlation shows that the information is then suitably formatted to guide behaviour. “Suitably formatted to guide behaviour” here means that the information is structured in such a way that the brain can apply a linear read out process to this representation to make a decision (importantly, this does not imply a causal link with behaviour). Carlson et al. (2014) demonstrated the promise of the distance-to-bound approach in a region of interest based analysis using fMRI. Here we go beyond this work by using the distance-to-bound method and a spatially unbiased fMRI-searchlight approach to create maps of where in the brain information can be used to guide behaviour.

## 2 Materials and Methods

In this study, we separately localised information that is decodable, and information that is suitably formatted to guide behaviour in the context of decodable information about visual objects and object categorisation behaviour. To ensure robustness and generality of our results, we analysed in parallel two independent fMRI datasets (Cichy et al., 2014, 2016), with different stimulus sets, and in relation to partly overlapping categorisation behaviours. Overall, this allowed us to investigate the relationship between decodable information from brain activity and categorisation behaviour for seven different distinctions: animate versus inanimate, faces versus bodies, human versus animal, natural versus artificial, tools versus not tools, food versus not food, and transport versus not transport. Note that the negative ‘not-X’ category was defined as all stimuli that did fall into one of the aforementioned classes. Categorisation reaction times for those stimuli were collected on Amazon’s Mechanical Turk. In this section, we describe the two-step searchlight procedure used to create decoding and correlation maps of areas involved in visual object categorisation.

### 2.1 Experimental design

#### Stimuli

Stimuli for experiment 1 consisted of 92 visual objects, segmented on a white background (Figure 1A). Stimuli consisted of animate and inanimate objects. The animate objects could be further divided into faces, bodies, humans and animals. Inanimate objects consisted of natural (e.g., plants or fruits) and man-made items (e.g., tools or houses). The stimulus set for experiment 2 consisted of 118 visual objects on natural backgrounds (Figure 1C). A small proportion of the objects (27) were animate. The inanimate objects included subcategories such as tools, or food items. In both experiments, participants were presented with the visual object stimuli while performing an orthogonal task at fixation. Stimuli were displayed at 2.9° (Experiment 1) and 4.0° (Experiment 2) visual angle with 500 ms duration. Images were displayed (overlaid with a grey fixation cross) for 500 ms in random order.

**Figure 1.**
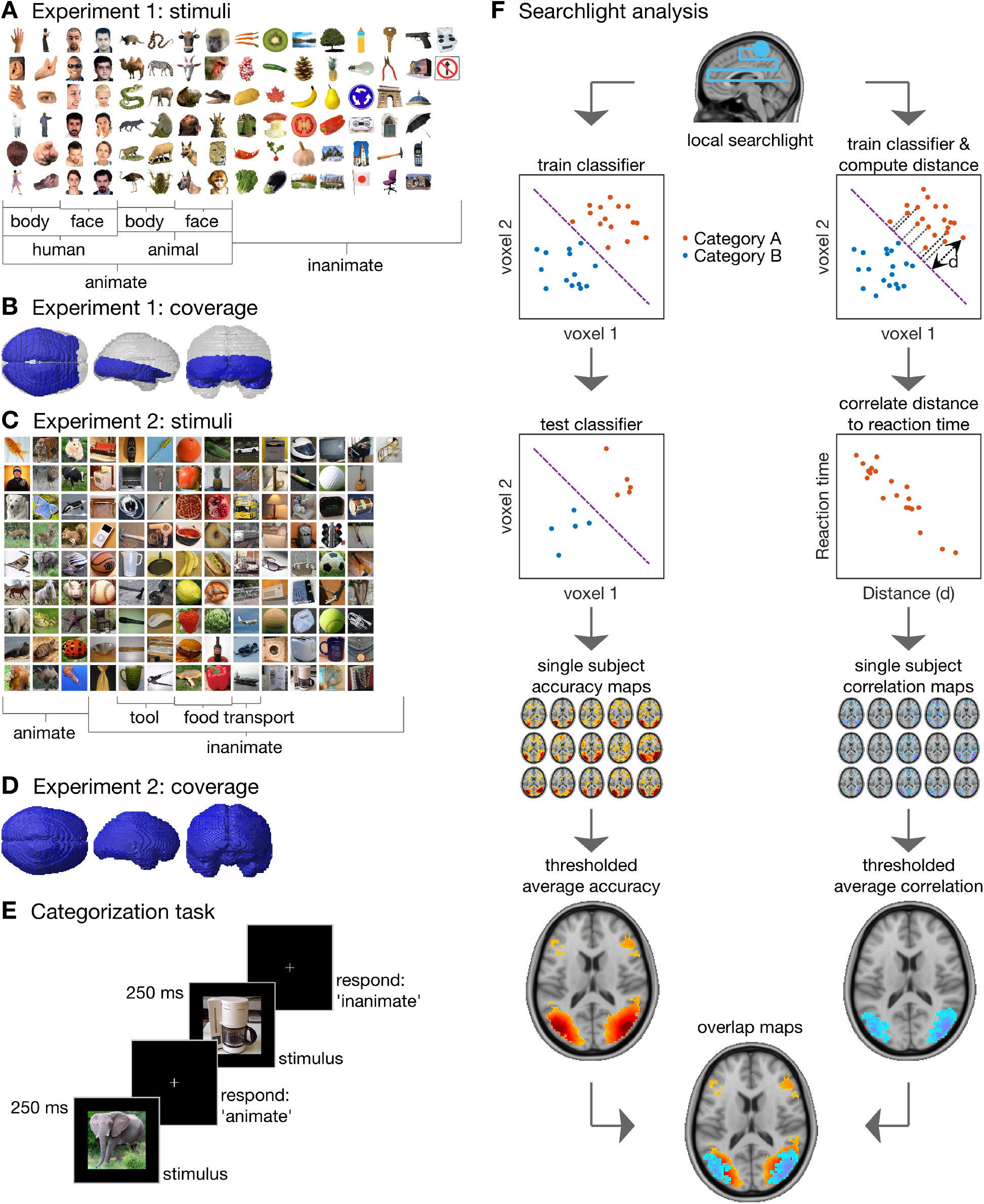
General experimental rationale. Stimuli (**A,C**) used to map fMRI brain responses and brain coverage (**B,C**) for fMRI study 1 and 2 respectively. **E.** Acquisition of reaction times on object categorisation tasks. Reaction times for categorisation contrasts were collected in a different pool of participants than the ones participating in the fMRI experiment. On each trial, a stimulus was displayed for 250ms, and participants categorised it into two categories (exemplarily here: animate vs inanimate) by pressing one of two keys. **F.** The two-partite approach to separately localize decodable information and information that is suitable for read out in behaviour. For both parts, a local cluster of neighbouring voxels (i.e., searchlight) was used to train a linear support vector machine (SVM) on an image category classification task (e.g., animacy). To localize decodable information, the classifier was tested on left-out data, storing the classification accuracy at the centre voxel of the searchlight. To localise information that was suitably formatted for read-out in a categorisation task, the distances of objects to the classifier hyperplane were correlated with the reaction times for the same object images on the same classification task. Repeated for every voxel, this resulted for each subject in one map of decoding accuracies and one of correlations. For visualisation, significant correlation voxels were superimposed on significant decoding accuracy voxels, each showing group average values in significant voxels.

#### fMRI recordings

The first experiment (Cichy et al., 2014) had high resolution fMRI coverage of the ventral visual stream (Figure 1B) from 15 participants with a 2 mm isotropic voxel resolution. The second experiment (Cichy et al., 2016) had whole brain from 15 participants with a 3 mm isotropic voxel resolution. In both experiments, at the start of a session, structural images were obtained using a standard T_1_-weighted sequence. fMRI data were aligned and coregistered to the T1 structural image, and then normalized to a standard MNI template. General linear models were used to compute t-values for each stimulus (92 and 118, respectively) against baseline.

#### Reaction time data

We obtained reaction times for the stimuli in multiple different categorisation contrasts (Figure 1A&B). For experiment 1, these were animate versus inanimate, face versus body, human versus animal, and natural versus artificial. For experiment 2, we tested animate versus inanimate, tool versus not tool, food versus not food, and transport versus not transport. The RTs were collected using Amazons Mechanical Turk (MTurk). For each of the categorisation contrasts, 50 unique participants performed a categorisation task using the same stimuli as were used in collecting the fMRI data. Participants were instructed to “Categorise the images as fast and accurate as possible using the following keys: (z for X, m for Y)”, where X and Y would be replaced with the relevant categories (e.g., animate and inanimate) for the contrast. On each trial, an image was presented for 500ms, followed by a black screen until the participant’s response (Figure 1C). The presentation order of the stimuli was randomized and stimuli did not repeat. This resulted in 50 reaction time values per exemplar (one for each participant). Each participant’s reaction times were z-scored. Next, we computed the median reaction time (across participants) for each exemplar. his resulted in one reaction time value per exemplar, which were used in the rest of the study.

### 2.2 Statistical Analysis

#### Searchlight procedure

For each categorisation contrast and subject, we used a searchlight approach (Haynes et al., 2007; Kriegeskorte et al., 2006) to create maps of decoding accuracy and of correlations between distance to the classifier boundary and categorisation reaction time. In contrast to pre-defined ROI’s, which are used to test a-priori hypotheses about the spatial origin of information in the brain, the searchlight results in a spatially unbiased map of decodable information. An overview of the approach is presented in Figure 1D.

To create the decoding accuracy maps, we used a standard searchlight decoding approach (Grootswagers, Wardle, & Carlson, 2017; Haynes, 2015; Kriegeskorte et al., 2006; Pereira, Mitchell, & Botvinick, 2009), as implemented in the CoSMoMVPA decoding toolbox (Oosterhof, Connolly, & Haxby, 2016). In detail, at each spatial location (voxel) in an fMRI image, a support vector machine (SVM) was used to classify visual object category based on local brain patterns, resulting in a map of classification accuracies. We then determined the subset of the locations at which brain patterns were suitably formatted for read-out by the brain using the distance-to-bound approach (Ritchie & Carlson, 2016) in a second searchlight analysis. Analogous to the decoding analysis, at each voxel, an SVM was trained to classify visual objects. Diverging at this point from the decoding approach we did not test the classifier, but rather obtained the distance for each exemplar to the hyperplane set by the SVM. We then correlated those distances to reaction times acquired in separate categorisation tasks. The contribution of each category was assessed individually, by performing the correlations separately for the two sides of the categorisation (e.g., one correlation for animate and one for inanimate exemplars). For each categorisation task this resulted in two correlation maps per subject. The maps of decoding accuracy and correlations were assessed for significance at the group level using sign-rank tests for random-effects inference. The results were thresholded at p < 0.05, using the false discovery rate (FDR; (Benjamini & Hochberg, 1995)) to correct for multiple comparisons at the voxel level.

#### Relating the results to topographical locations of the visual system

For the animacy categorisation contrasts, we identified the locations of the significant voxels with respect to ROIs of the visual system. The significant voxels in the decoding maps and correlation maps were compared to probabilistic topographic maps of visual processing areas (Wang, Mruczek, Arcaro, & Kastner, 2015), which represent for each voxel the visual area with the highest probability. A percentage score for each ROI was then computed, reflecting the percentage of voxels in this ROI that were significant at the group level. We obtained a bootstrapped distribution of percentage scores for each ROI by repeating this procedure 10,000 times, while randomly sampling the subjects with replacement and recomputing the group level statistics. We report the 5^th^, 50^th^ and 95^th^ percentiles of this distribution. This approach allows quantifying the difference between the number of decoding voxels and correlation voxels per visual ROI.

## 3 Results

We examined the relationship between decodable information and information that is suitably formatted for read-out by the brain in the context of decodable information about visual objects and object categorisation behaviour. We determined the relationship between decodable information and behaviour separately. First, we determined where information about objects is present in brain patterns using decoding in a standard fMRI searchlight decoding analysis (Haynes et al., 2007; Kriegeskorte et al., 2006). We then determined the subset of the locations at which brain patterns were suitably formatted for read-out by the brain using the distance-to-bound approach (Ritchie & Carlson, 2016) in a second searchlight analysis. The subject-specific searchlight results were subjected to inference statistics at the group level using one-sided sign rank tests and thresholded at p < 0.05 (fdr-corrected for multiple comparisons across voxels).

### 3.1 A subset of locations that have decodable information about animacy also had information suitably formatted for animacy categorisation behaviour

Animacy is a pervasive and basic object property according to which any object can be classified as animate or inanimate (Caramazza & Shelton, 1998). Previous studies have shown that the division of animate versus inanimate objects is reflected in the large-scale architecture of high-level visual areas such as the ventral temporal cortex (VTC) (Caramazza & Shelton, 1998; Grill-Spector & Weiner, 2014; Kriegeskorte et al., 2008), However, it has also been shown that animacy can be decoded not only from VTC, but from the whole ventral visual stream (Cichy et al., 2016; Grill-Spector & Weiner, 2014; Long, Yu, & Konkle, 2017). Furthermore, categorical object responses have also been found in the dorsal visual stream (Bracci, Daniels, & op de Beeck, 2017; Freedman & Assad, 2006; Konen & Kastner, 2008) and in frontal areas (Freedman, Riesenhuber, Poggio, & Miller, 2001, 2003). This prompts the question of where in the visual system object representations are suitably formatted for read-out by the brain for animacy decisions.

Corroborating previous studies, we found decodable information about animacy in the entire ventral visual stream from the occipital pole to anterior ventral temporal cortex (Figure 2AB, Table 1AE, N = 15, one-sided sign-rank test, p < 0.05 fdr-corrected). In addition, we found decodable information in dorsal and prefrontal cortex (Figure 2B) in experiment 2 which had full brain coverage. Localising the brain representations suitable to guide animacy categorisation behaviour (using the distance-to-bound approach) revealed convergent evidence across experiments that only a subset of voxels containing decodable information fulfilled this criterion. In detail, distance-RT-correlations for animate objects were strongest in the high-level regions of the ventral and the dorsal stream. For inanimate objects, we found no voxels with significant distance-RT-correlations (Carlson et al., 2014; Grootswagers, Ritchie, Wardle, Heathcote, & Carlson, 2017).

**Figure 2.**
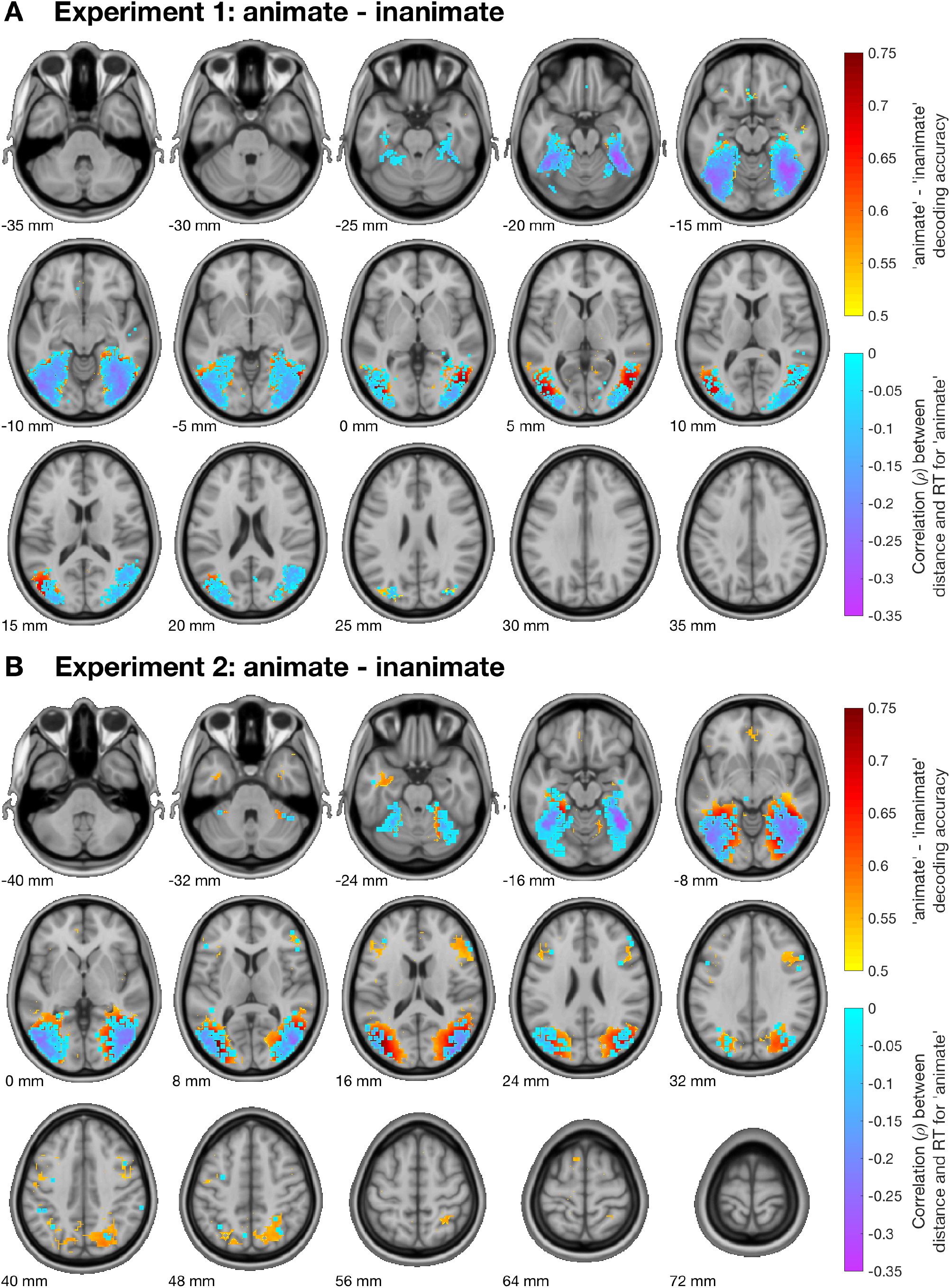
Relationship between decodable information and categorisation behaviour for animacy. Decodable information is shown in hot colours and distance-RT-correlations in cool colours. Colour intensities reflect the mean across subjects. Only significant voxels (N=15, sign-rank test, p<0.05 fdr-corrected) are shown. Data are projected onto axial slices of a standard T_1_ image in MNI space. **A.** In experiment 1, decodable animacy information (hot colours) was found throughout the ventral stream. A correlation between distance to the classifier boundary and reaction time for animate stimuli (cool colours) was found in a subset of these areas. The colour intensities depict the mean across subjects. **B.** The results of the analysis for experiment 2 corroborated these findings, and showed decodable information in prefrontal areas and in the dorsal visual stream. Correlations between distance and reaction time were also present in the dorsal stream.

**Table 1.**
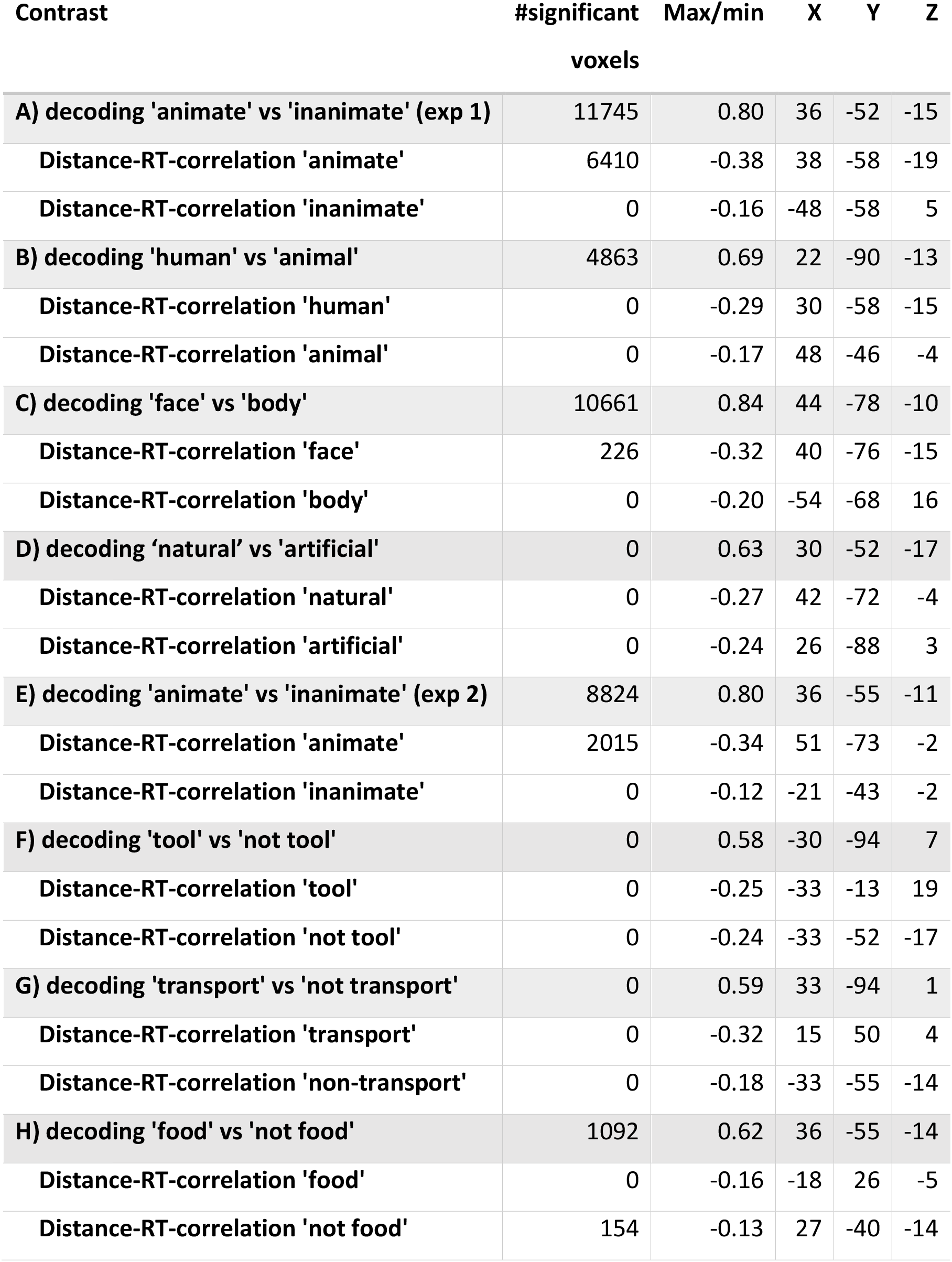
Results for all categorisation contrasts. For all categorisation contrasts, we report the number of significant voxels (after correction for multiple comparisons), its peak value (maximum for decoding or minimum for distance-RT-correlation), and the peak’s location in MNI-XYZ coordinates.

| Contrast | #significant voxels | Max/min | X | Y | Z |
| --- | --- | --- | --- | --- | --- |
| A) decoding 'animate' vs 'inanimate' (exp 1) | 11745 | 0.80 | 36 | -52 | -15 |
| Distance-RT-correlation 'animate' | 6410 | -0.38 | 38 | -58 | -19 |
| Distance-RT-correlation 'inanimate' | 0 | -0.16 | -48 | -58 | 5 |
| B) decoding 'human' vs 'animal' | 4863 | 0.69 | 22 | -90 | -13 |
| Distance-RT-correlation 'human' | 0 | -0.29 | 30 | -58 | -15 |
| Distance-RT-correlation 'animal' | 0 | -0.17 | 48 | -46 | -4 |
| C) decoding 'face' vs 'body' | 10661 | 0.84 | 44 | -78 | -10 |
| Distance-RT-correlation 'face' | 226 | -0.32 | 40 | -76 | -15 |
| Distance-RT-correlation 'body' | 0 | -0.20 | -54 | -68 | 16 |
| D) decoding 'natural' vs 'artificial' | 0 | 0.63 | 30 | -52 | -17 |
| Distance-RT-correlation 'natural' | 0 | -0.27 | 42 | -72 | -4 |
| Distance-RT-correlation 'artificial' | 0 | -0.24 | 26 | -88 | 3 |
| E) decoding 'animate' vs 'inanimate' (exp 2) | 8824 | 0.80 | 36 | -55 | -11 |
| Distance-RT-correlation 'animate' | 2015 | -0.34 | 51 | -73 | -2 |
| Distance-RT-correlation 'inanimate' | 0 | -0.12 | -21 | -43 | -2 |
| F) decoding 'tool' vs 'not tool' | 0 | 0.58 | -30 | -94 | 7 |
| Distance-RT-correlation 'tool' | 0 | -0.25 | -33 | -13 | 19 |
| Distance-RT-correlation 'not tool' | 0 | -0.24 | -33 | -52 | -17 |
| G) decoding 'transport' vs 'not transport' | 0 | 0.59 | 33 | -94 | 1 |
| Distance-RT-correlation 'transport' | 0 | -0.32 | 15 | 50 | 4 |
| Distance-RT-correlation 'non-transport' | 0 | -0.18 | -33 | -55 | -14 |
| H) decoding 'food' vs 'not food' | 1092 | 0.62 | 36 | -55 | -14 |
| Distance-RT-correlation 'food' | 0 | -0.16 | -18 | 26 | -5 |
| Distance-RT-correlation 'not food' | 154 | -0.13 | 27 | -40 | -14 |

### 3.2 The proportion of region-specific representations suitably formatted for behaviour increases along the ventral stream and decreases along the dorsal stream

We next explicitly determined the degree to which representations in single brain regions within the ventral and dorsal streams are suitably formatted for behaviour. For this we parcellated the cortex (Figure 3A) using a probabilistic topographic map of visual processing areas (Wang et al., 2015). For each region, we calculated the ratio between the number of significant voxels in the decoding analysis and the total number of voxels, so that a high ratio indicates that a large part of a region contains object representations with categorical information. Similarly, we calculated the ratio between the number of significant voxels in the distance-to-bound analysis and the total number of voxels. Here, a high ratio indicates that a large part of a region contains object representations that are suitably formatted for read out in a categorisation task.

**Figure 3.**
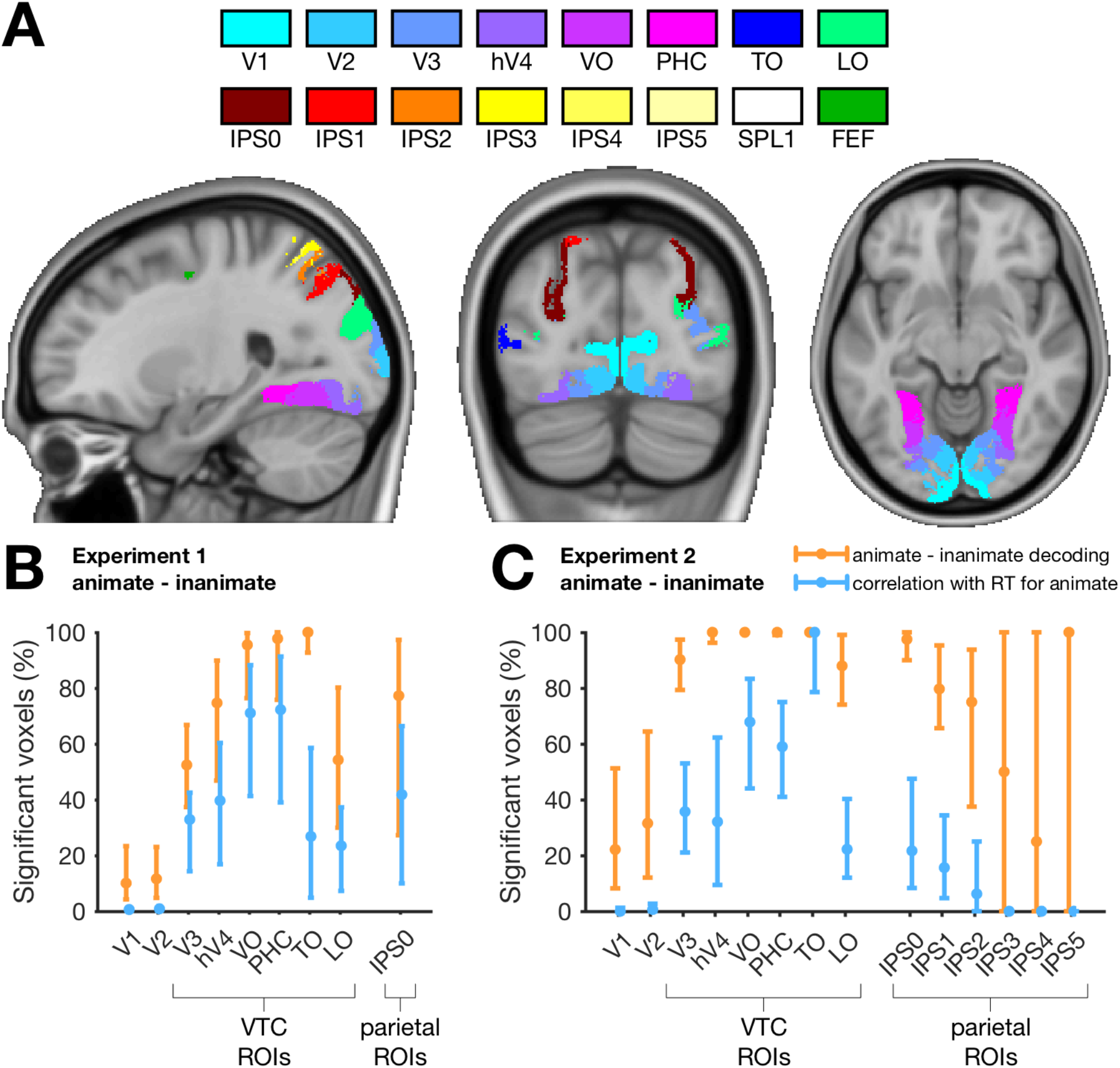
Quantifying the decodable information in visual areas and their contribution to categorisation behaviour. **A.** Locations of topographical ROIs of the visual system (Wang et al., 2015), containing early visual cortex (EVC) areas V1 and V2, mid-level areas V3 and hV4, high level ventral occipital (VO) and parahippocampal cortex (PHC), temporal occipital (TO) and lateral occipital (LO) areas, areas in the intra-parietal sulcus (IPS), the superior parietal lobule (SPL), and the frontal eye fields (FEF). **B-C.** The ratio between significant voxels in an ROI and the size of the ROI. Orange points show the ratio of voxels within the ROI that had significant animacy decoding performance. Blue points show the ratio of voxels with a significant correlation between distance to the hyperplane and RT for ‘animate’. The lower, middle and upper points on these lines indicate 5^th^, 50^th^, and 95^th^ percentiles (bootstrapping of participants 10,000 times). These results quantify the increasing contribution of early to late areas in the ventral visual stream to animacy categorisation behaviour.

In the ventral stream, our results suggest that these ratios increase with processing stage, from early visual areas to high-level visual areas, with highest ratios in ventral occipital (VO) and parahippocampal (PHC) cortex (Figure 3 B&C). In contrast, in the dorsal stream we observed a decrease of the correlation ratio with processing stage. In addition, significant animacy decoding information was found in similar proportions in the ventral-temporal areas as in lateral-occipital areas, however, the proportion of voxels with information suitable for categorisation was lower in lateral-occipital areas. This is consistent with the notion that while both these regions contain object representations, the VTC contains location-invariant representations which are essential for object categorisation (Cichy et al., 2013; Haushofer et al., 2008; Schwarzlose, Swisher, Dang, & Kanwisher, 2008; Williams et al., 2007). The results were similar between experiments, with the exception for area TO, which had a smaller proportion of voxels with RT-correlations in experiment 1. It is possible that this difference was caused by the differences between the stimuli (e.g., segmented objects versus objects in scenes) used in the experiments. Alternatively, this difference could be attributed to the size of the searchlight sphere, which was larger in experiment 2 than in experiment 1 due to their different voxel sizes.

In sum, these results show that representations along the ventral stream are suitably formatted for read-out of categorical information (Cichy et al., 2013; Grill-Spector & Weiner, 2014). In contrast, representations in the dorsal stream might be shaped for the read-out in different tasks (Bracci et al., 2017; Freud et al., 2017). These results also suggest that intermediate stages along the ventral and dorsal streams may be similar or partly shared, as suggested by the similar ratios of information suitable for read-out.

### 3.3 Decodable information about subordinate categorisation tasks is also suitably formatted for categorisation behaviour

While animacy categorisation may be based on large-scale representational differences in the visual brain (Carlson, Tovar, Alink, & Kriegeskorte, 2013; Downing, Chan, Peelen, Dodds, & Kanwisher, 2006; Grill-Spector & Weiner, 2014; Kriegeskorte et al., 2008), subordinate categorisation tasks (e.g., faces, bodies, tools) may depend more on fine grained patterns in focal brain regions (Downing, Jiang, Shuman, & Kanwisher, 2001; Downing & Peelen, 2016; Kanwisher, McDermott, & Chun, 1997). Here, we tested whether decodable information about subordinate category membership is also suitably formatted for read out in respective categorisation tasks. We tested two subordinate contrasts for the animate exemplars in experiment 1: face versus body, and human versus animal using the same general procedure as for animacy. We found that both contrasts were decodable (Table 1B-C). We found a significant correlation between distance to the classifier hyperplane and reaction times for faces in the face versus body task (Figure 4A). We found no significantly decodable information or significant correlations for the natural versus artificial objects (Table 1D). Of the subordinate categorisation contrasts in experiment 2 (food, transport or tool versus everything else), transport and tool versus everything else were not significantly decodable information nor had they significant correlations (Table 1F-G). Food versus not food resulted in significant decodable information, and significant distance-RT correlations were present for this contrast in the ‘not food’ category (Figure 4B, Table 1H). Taken together, for some subordinate categorisation contrasts that were decodable, we were successful in localising brain patterns suitably formatted for read-out in behaviour.

**Figure 4.**
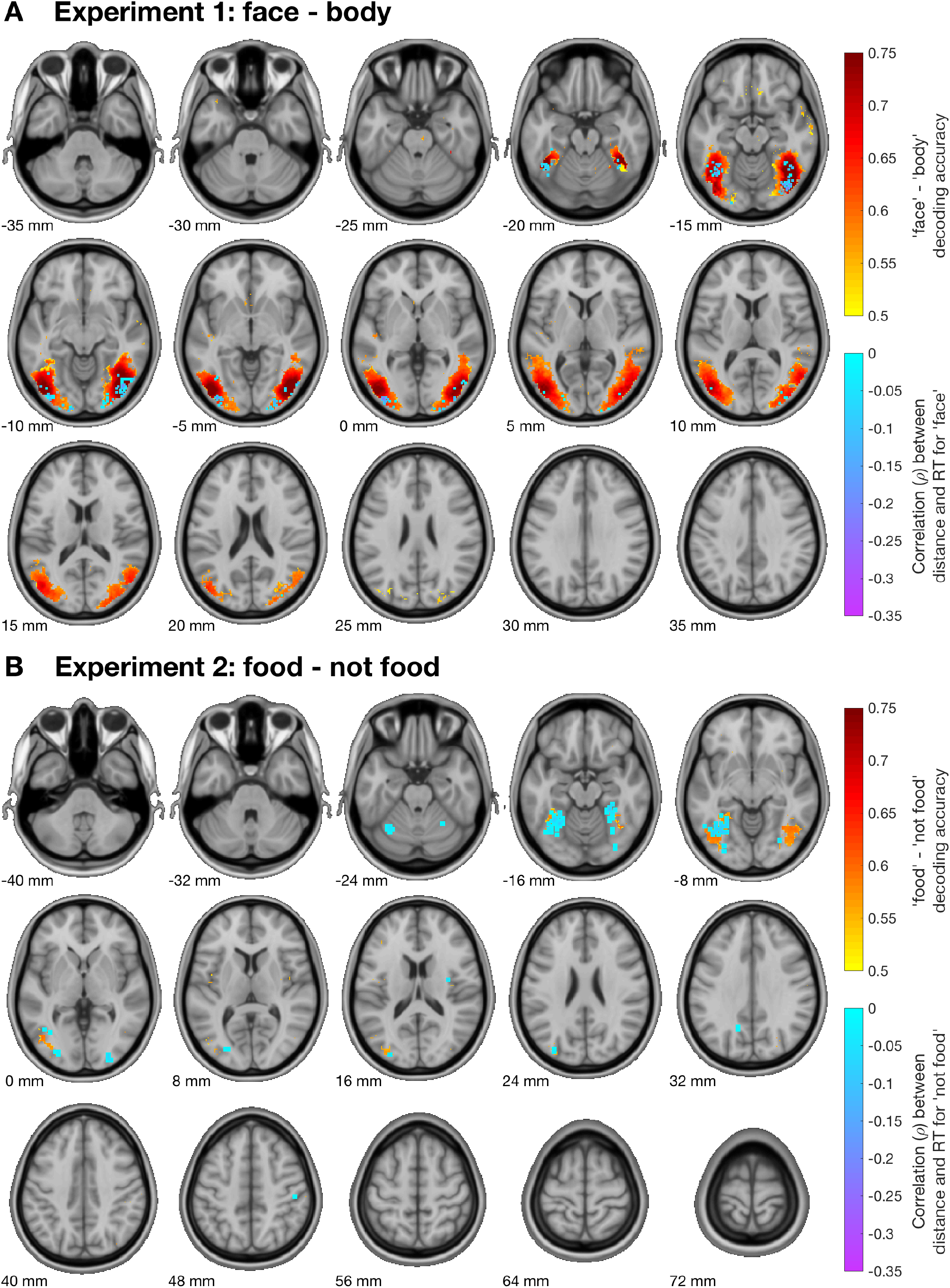
Relationship between decodable information and behaviour for subordinate categorisation tasks. Decodable information is shown in hot colours and distance-RT-correlations in cool colours. Colour intensities reflect the mean across subjects. Only significant voxels (N=15, sign-rank test, p<0.05 fdr-corrected) are shown. Data are projected onto axial slices of a standard T_1_ image in MNI space. **A.** In experiment 1, decodable face versus body information (hot colours) was found in the entire ventral stream. A distance-RT-correlation for the face stimuli (cool colours) was found in a subset of these areas. **B.** In experiment 2, food versus not food was decodable in some areas in the ventral visual stream. A distance-RT-correlation for the ‘not food’ stimuli was found in a subset of these areas.

## 4 Discussion

### 4.1 Dissociating between decodable information and information that is used in behaviour

The aim of this study was to examine where in the brain decodable information is suitably formatted for read-out by the brain in behaviour. We found that only a subset of information that is decodable could be related to behaviour using the distance-to-bound approach, which argues for a partial dissociation between decodable information and information that is relevant for behaviour. This speaks to a current challenge in neuroimaging, which is to show that information visible to the experimenter is in fact used by the brain (de-Wit et al., 2016; Ritchie et al., 2017). To illustrate, consider the question about what regions are used by the brain to perform an object animacy categorisation task (DiCarlo, Zoccolan, & Rust, 2012; Grill-Spector & Weiner, 2014). On its own, the result of the animacy decoding searchlight might be interpreted as the brain using animacy information from anywhere in the ventral stream. However, when investigating this interpretation directly using the distance-RT-correlation results, it becomes clear that object animacy information is suitably represented for read-out in mid- and high-level visual areas only.

It is important to note that not finding a correlation between distance to the classifier hyperplane and RT does not imply that the information revealed using the decoding approach is irrelevant or epiphenomenal. The distance-to-bound approach taken here makes specific assumptions about the brain’s read-out process, such as distance in representational space as the measure for evidence, and a monotonic relationship between distance and reaction time (Ritchie & Carlson, 2016). Note that this model of readout follows from the assumptions behind the decoding methods (Ritchie & Carlson, 2016; Ritchie et al., 2017). While the model may not be perfect, our results stress the importance of explicitly testing models of readout when decoding information from the brain. Finding the correct model of readout would significantly increase the capacity of cognitive neuroscience to infer brain-behaviour relationships. Other assumptions follow from those imposed by the decoding approach, such as the binary classification, the size of the searchlight radius, the choice of classifier. For example, it could be that the representations are relevant in a different task (Grootswagers, Ritchie, et al., 2017; Ritchie & Carlson, 2016), or that read-out involves pooling over larger spatial scales or multiple brain areas. Therefore, the current approach only allows the positive inference on the level of suitability of decoded information for behaviour in the context of the current task and decoding parameters. On the other hand, a correlation with behaviour still does not prove that the information is used by the brain, but it shows that the information is at least formatted in a way that is suitable to be used by the brain for decisions. Future work can use causal measures (e.g., TMS) targeting the areas highlighted in the current results.

### 4.2 The contribution of ventral and dorsal visual regions to categorisation behaviour

We found that neural representations suitably formatted for behaviour in categorisation were most prominently located in the anterior regions of the VTC. This corroborates previous studies (Afraz, Kiani, & Esteky, 2006; Carlson et al., 2014; Hong, Yamins, Majaj, & DiCarlo, 2016; Hung, Kreiman, Poggio, & DiCarlo, 2005), and reinforces the tight link between VTC and visual categorisation behaviour. In these areas, our results provide converging evidence for the (implicit) assumption made in neuroimaging studies, which is that information that is available to the experimenter is also available for read out by the brain in behaviour (cf. de-Wit et al., 2016).

However, we found that correlations between distance to boundary and RT were not restricted to anterior regions of the VTC, but were also prominent in V3 and hV4. This is consistent with the view that lower level visual features encoded in mid-level visual regions could aid faster read-out of category information. V4 is thought of as an intermediate stage of visual processing that aggregates lower level visual features into invariant representations (Riesenhuber & Poggio, 1999). It has been proposed that direct pathways from V4 to decision areas allow the brain to exploit visual feature cues for fast responses to ecologically important stimuli (Hong et al., 2016; Kirchner & Thorpe, 2006; Thorpe, Fize, & Marlot, 1996), such as identifying faces (Crouzet, Kirchner, & Thorpe, 2010; Honey, Kirchner, & VanRullen, 2008). An alternative possibility is that read out is not happening directly from V4, but its representational structure is shaped by the low-level feature differences in animacy. This structure is then largely preserved when it is communicated to more anterior areas, leading to similar distance-RT-correlations. Both of these accounts are also consistent with recent findings that show differential responses for object categories in mid-level visual areas (Long et al., 2017; Proklova et al., 2016). The extent to which visual features contribute to the read-out process could be further investigated by using the approach from this study with different stimulus sets that control for these features (Kaiser, Azzalini, & Peelen, 2016; Long et al., 2017; Proklova et al., 2016).

We found that distance-RT-correlations were also present in early parietal areas. The classical view is that the ventral and dorsal visual streams are recruited for different function (Ungerleider & Mishkin, 1982). However, areas in the ventral and dorsal streams have been found to exhibit similar object-selective responses (Freud et al., 2017; Konen & Kastner, 2008; Sereno & Maunsell, 1998; Silver & Kastner, 2009). Consistent with this, we found similar RT-distance-correlations in mid-level areas in the ventral and dorsal streams. However, our results also showed that the proportion of correlations decreased along the dorsal stream, while they increased along the ventral stream. This suggests that representations in the ventral and dorsal streams undergo similar transformations at first, and then diverge for different goals.

### 4.3 Without a task, neural object representations in the VTC are formatted for read-out in categorisation decisions

Here, the fMRI participants performed an orthogonal task, and were not actively categorising. Despite this, categorisation reaction times could still be predicted from representations in the visual stream. This highlights that, without a categorisation task, information in the visual system is represented in a way that is suitable for read out in behaviour (Carlson et al., 2014; Ritchie, Tovar, & Carlson, 2015). This representation possibly reflects a more general property of the object that aids its categorisation, such as how typical it is for that category (Grootswagers, Ritchie, et al., 2017; Iordan, Greene, Beck, & Fei-Fei, 2016), or how frequently we encounter the object in our lives. In addition, the orthogonal task in the scanner has the advantage that it avoids RT- and difficulty confounds (see e.g., Hebart & Baker, 2017; Woolgar, Golland, & Bode, 2014). Future studies might use the distance-to-bound approach with participants actively performing the same task in the scanner, where we predict that areas involved in the decision making and execution processes would contain information that correlates with reaction times. For example, some areas preferentially represent task-relevant information, such as areas in the prefrontal cortex (Duncan, 2001; Jackson, Rich, Williams, & Woolgar, 2016; Woolgar, Jackson, & Duncan, 2016), and in the parietal stream (Bracci et al., 2017; Freedman & Assad, 2016; Jeong & Xu, 2016). In the absence of an animacy categorisation task, one would predict that animacy information would not be strongly represented in these areas. Yet, our results showed that animacy information can be decoded from prefrontal and parietal areas when participants perform an orthogonal task. However, our results did not provide evidence that the animacy information in these areas was suitably formatted for readout. This again argues for a dissociation between information that can be decoded, and information that is suitable for read out in behaviour. A prediction that follows from this is that performing an active object categorisation task in the scanner would change the representations in these task-relevant areas so that they become predictive of reaction times (Bugatus, Weiner, & Grill-Spector, 2017; McKee, Riesenhuber, Miller, & Freedman, 2014). Similarly, representations can change when participants perform different tasks on the same stimuli, such as categorising a specific feature (e.g., colour), for which suitably formatted information would be predicted in other areas.

### 4.4 Asymmetric distance-RT-Correlations in binary categorisation tasks

In both experiments, we found correlations between distance and reaction times for animate stimuli, but none for the inanimate stimuli. This is consistent with previous work (Carlson et al., 2014; Grootswagers, Ritchie, et al., 2017; Ritchie et al., 2015), which argued that this discrepancy might be caused by inanimate being a negatively defined category (i.e., “not animate”). Under this hypothesis the animacy categorisation task can be performed by collecting evidence for animate stimuli and responding inanimate only when not enough evidence was accumulated after a certain amount of time. Here, we tested a prediction of this hypothesis by contrasting two positively defined categories, face versus body, and found that there was a distance-RT-correlation only for faces. This goes against the notion of the negative definition of inanimate as the main reason for a lack of correlation. However, it still is possible that observers still treated these tasks as ‘A’ or ‘NOT A’, with ‘A’ being the category that is easiest to detect (Grootswagers, Ritchie, et al., 2017). For example, perceptual evidence for a face would be easier to obtain than evidence for a body-part, as faces share low level visual features (Crouzet & Thorpe, 2011; Honey et al., 2008; Wu, Crouzet, Thorpe, & Fabre-Thorpe, 2015). Thus, while not explicitly specified as a negative category, it could have been treated as such.

This suggests that the binary categorisation might be an unnatural way of approaching human categorisation behaviour in the real world. Other operationalisations such as picture naming or visual search may be better suited to capture the relevant behaviours (cf. Krakauer, Ghazanfar, Gomez-Marin, MacIver, & Poeppel, 2017). Still, it is important to note that the binary task matches the brain decoding task performed by the classifier. The above-chance decoding accuracy in the brain decoding task is commonly interpreted as a similar dichotomy in the brain’s representation that the brain can use in a decision. However, when only the information in one of the categories (i.e., animals or faces) can be used to predict decision behaviour, as shown here, then this interpretation needs to be revisited.

### 4.5 Limitations of the approach

Our results highlight the importance of relating decoding to behaviour and demonstrated one possible methodology to address this issue. However, the approach taken here is subject to a set of limitations which may preclude its application in other settings. Firstly, here we studied a binary visual object categorisation task. It is not possible to describe all behaviours as binary tasks, and reaction times are not always a meaningful measure for behaviour. This can restrict the generalisability of the current approach to other domains. Secondly, finding an RT-correlation does not reveal the source of the variance in evidence for a decision. As the method remains correlational, it is important to stress that it can only go as far to show that information is suitably formatted to be used by the brain for decisions, and that the critical test of this relationship will require causal measures. In the animacy task, one possible source of variance is typicality, which modulates animacy categorisation (Posner & Keele, 1968; E. H. Rosch, 1973; E. Rosch & Mervis, 1975) and decoding performance (Iordan et al., 2016), and typicality ratings have been shown to correlate with distance to the classifier decision boundary (Grootswagers, Ritchie, et al., 2017). Yet, there is always the possibility of other covariates. For example, larger or more colourful objects may be easier to identify and therefore have a faster reaction times and larger distances, and it is not always feasible to control for all possible covarying features. Note that this is also a strength of the approach; if colourful objects are indeed easier to identify (i.e., the brain is using the feature), then areas where this feature is represented would have stronger correlations between distance to boundary and reaction times. If a stimulus property is thought to (unintentionally) drive decoding and behaviour in the same way, then this property should be controlled for in the stimulus set.

### 4.6 Conclusion

In this study, we combined the distance-to-bound approach (Ritchie & Carlson, 2016) with a searchlight decoding analysis to find brain areas with decodable information that is suitable for read-out in behaviour. Our results showed that decodable information is not always equally suitable for read-out by the brain in behaviour. This speaks to the current debate in neuroimaging research about whether the information that we can decode is the same information that is used by the brain in behaviour (de-Wit et al., 2016).

## Acknowledgements

This research was supported by an Australian Research Council Future Fellowship (FT120100816) and an Australian Research Council Discovery project (DP160101300) awarded to T.A.C., and a German Research Foundation grant (CI-241/1-1) awarded to R.M.C. The authors acknowledge the University of Sydney HPC service for providing High Performance Computing resources. The authors declare no competing financial interests.

